## Supplementary figures for "A revamped rat reference genome improves the discovery of genetic diversity in laboratory rats"

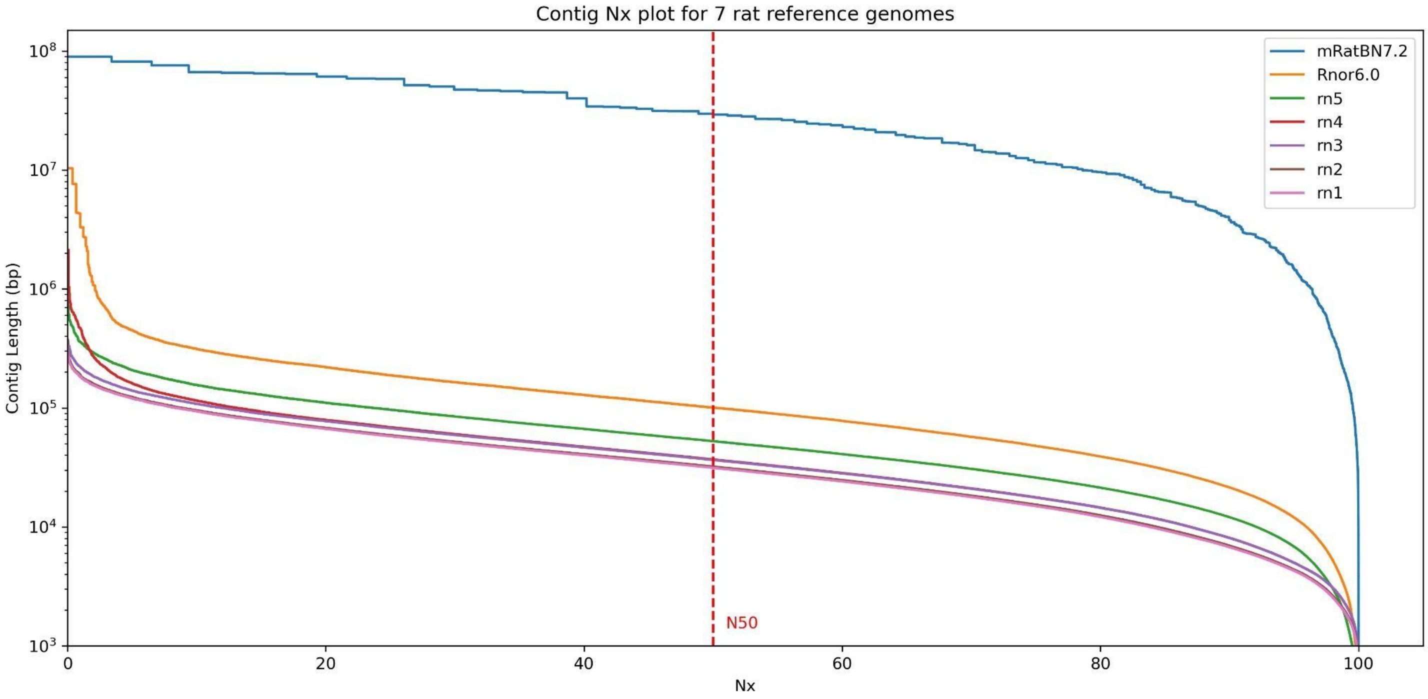


**Figure S1. The Contig Nx plot for mRatBN7.2 and 6 prior rat reference genomes.** The red dashed line indicates the contig N50 values. Small improvements in contig continuity are observed for the previous updates, with the most significant improvement (~290X) coming from mRatBN7.2

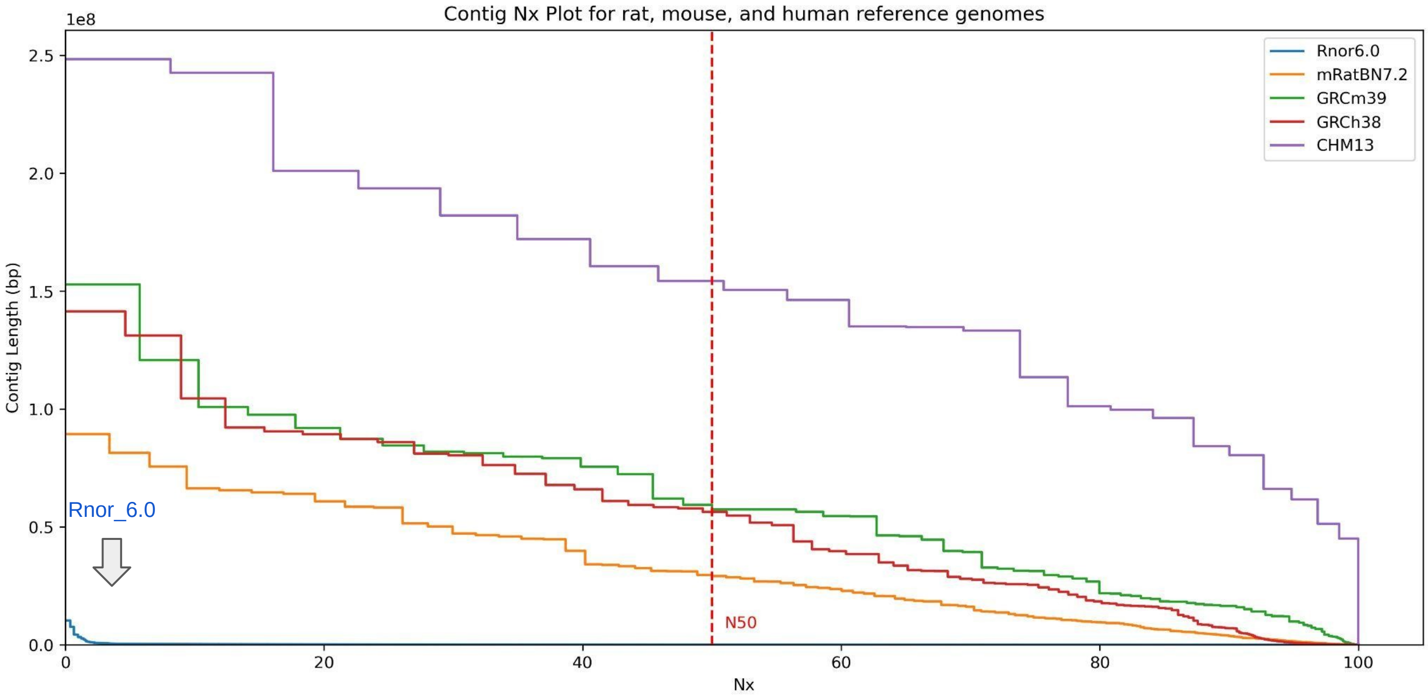


**Figure S2.The Contig Nx plot for the Rat, Mouse, and Human reference genomes.** The red dashed line indicates the contig N50 values. The Nx curve for Rnor6.0 is very low and partially overlapping with the X axis. The CHM13, being the first truly gapless human genome, is in its own tier in terms of assembly continuity. The GRCh38 released in 2014 and the GRCm39 released in 2020 have similar continuity. Although mRatBN7.2 is still lagging behind in continuity compared to the current human and mouse reference genomes, it represent a very significant improvement over the current rat reference genome Rnor_6.0 (Orange vs. Cyan line).

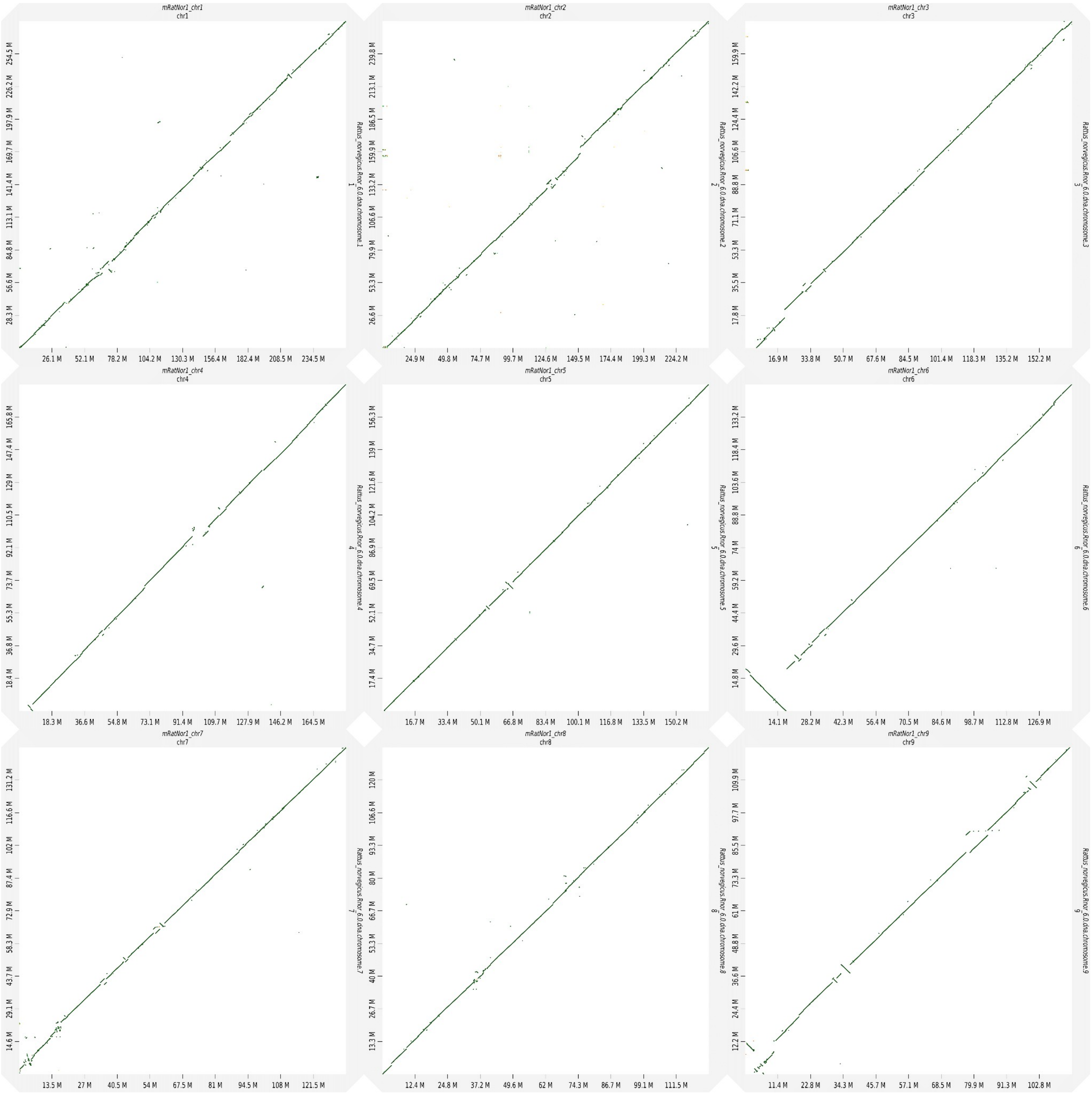


**Figure S3. Chromosomal dot plots between Rnor_6.0 and mRatBN7.2**. mRatBn7.2 is on the y-axis and Rnor_6.0 is on the x-axis.

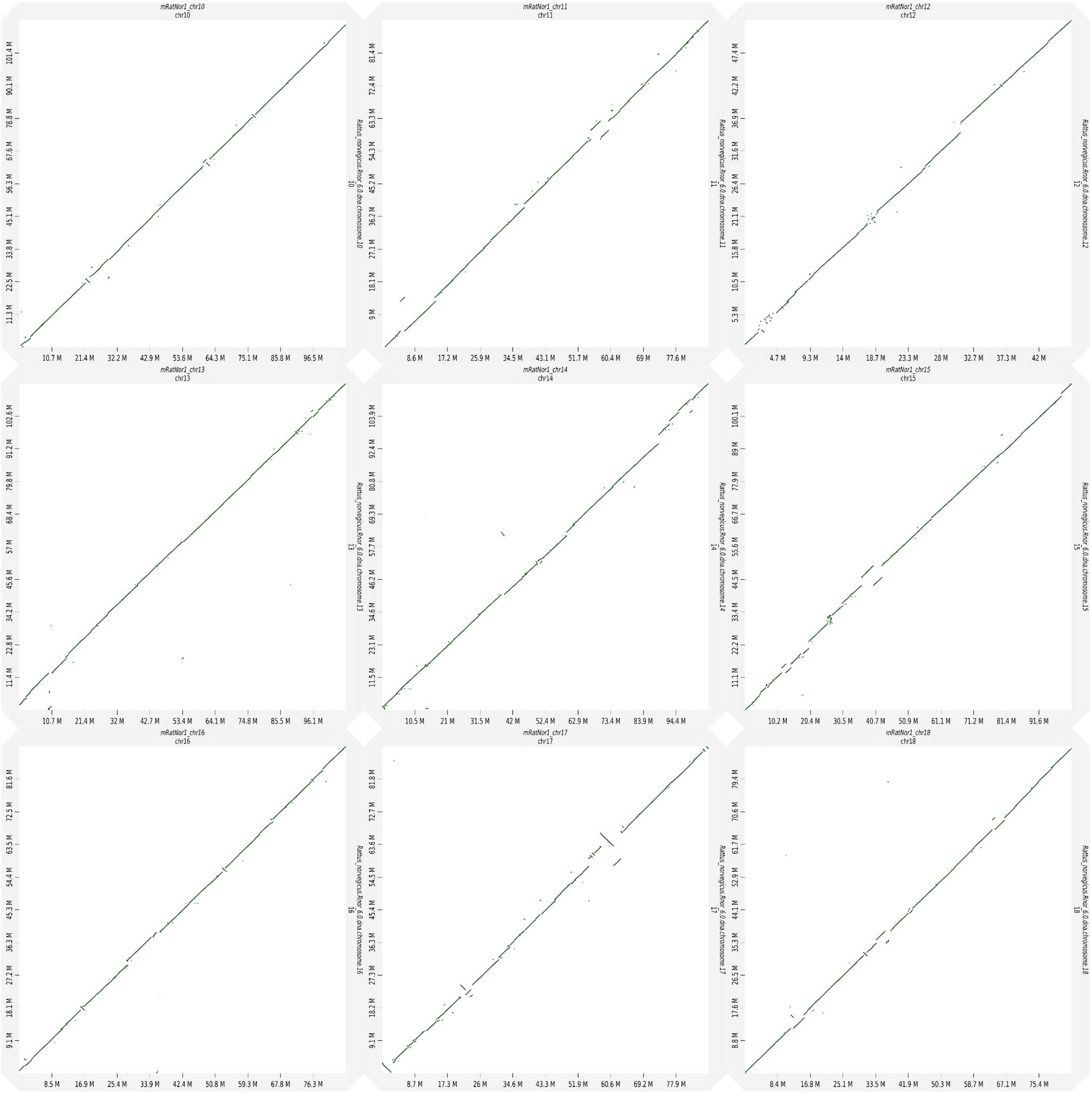

**Figure S3 Continued.**

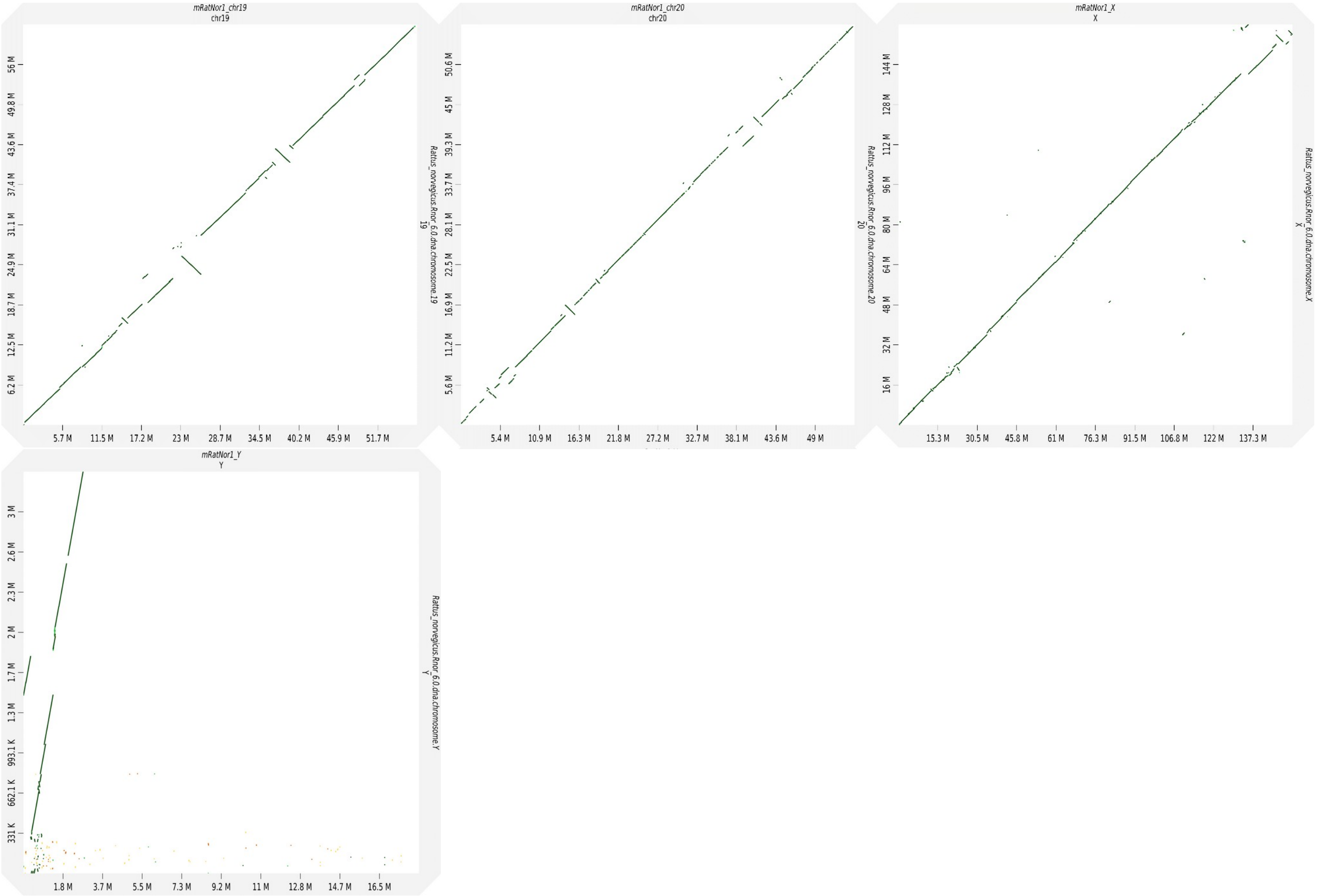


#### Figure S3 Continued

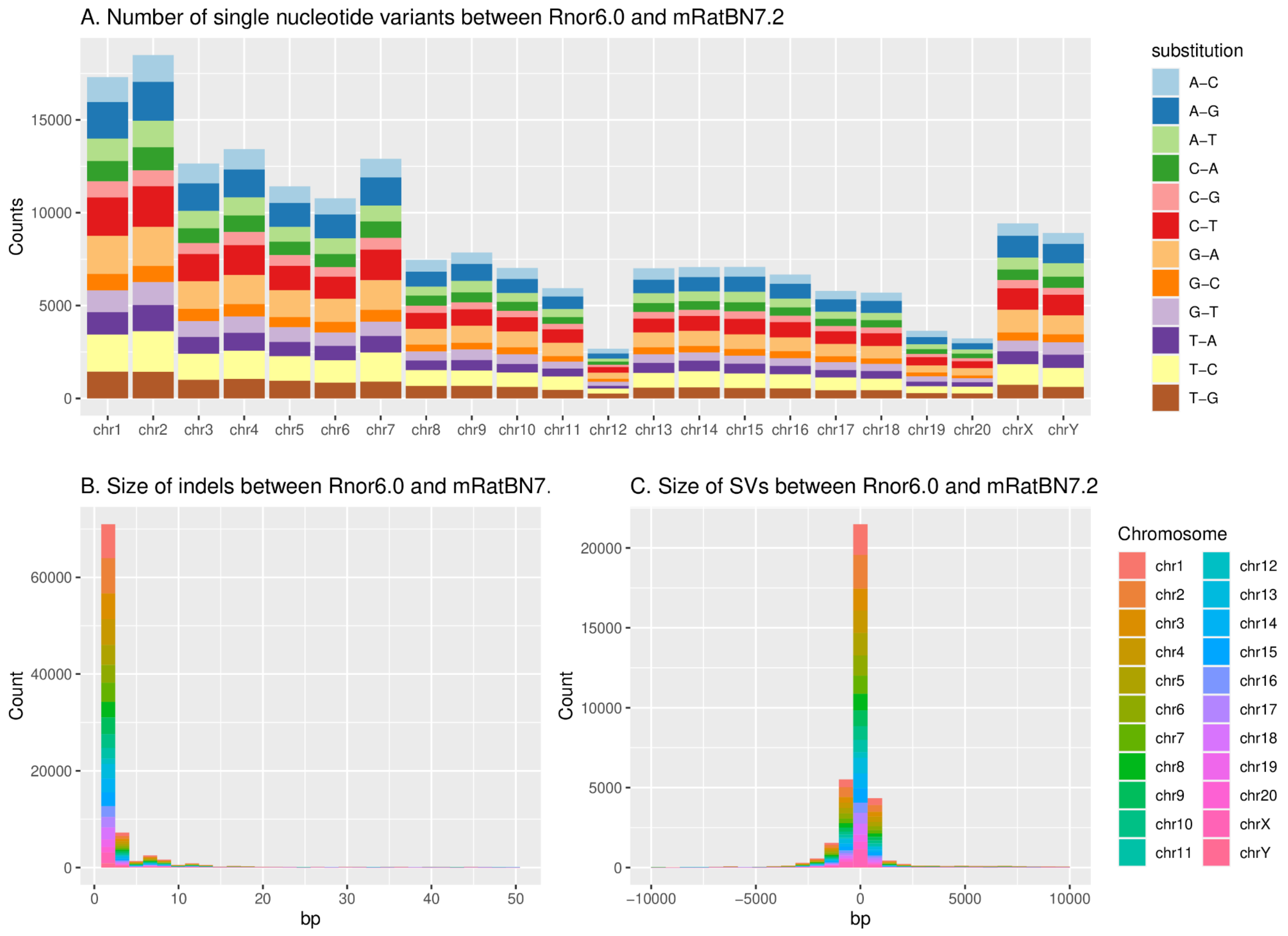


**Figure S4: SNP and structural variants between Rnor_6.0 and mRatBN7.2. A)** The substitution frequency per chromosome for Rnor6.0 and mRatBN7.2. **B)** The size of indels between Rnor 6.0 and mRatBN7.2. **C)** The size of SVs between Rnor6.0 and mRatBN7.2.

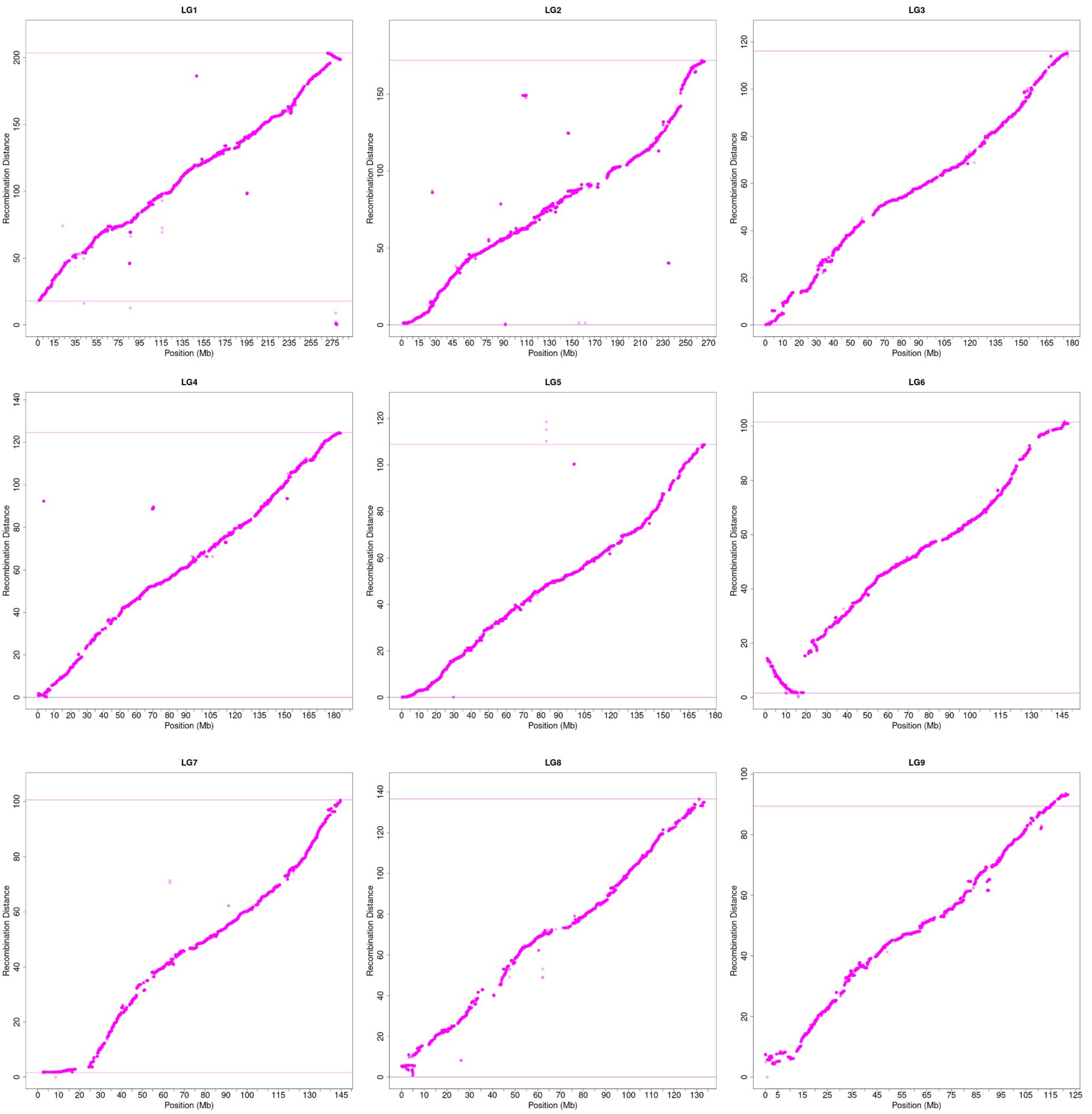


**Figure S5. The order of genetic markers and the distances from a rat genetic map compared to their locations in Rnor_6.0.**

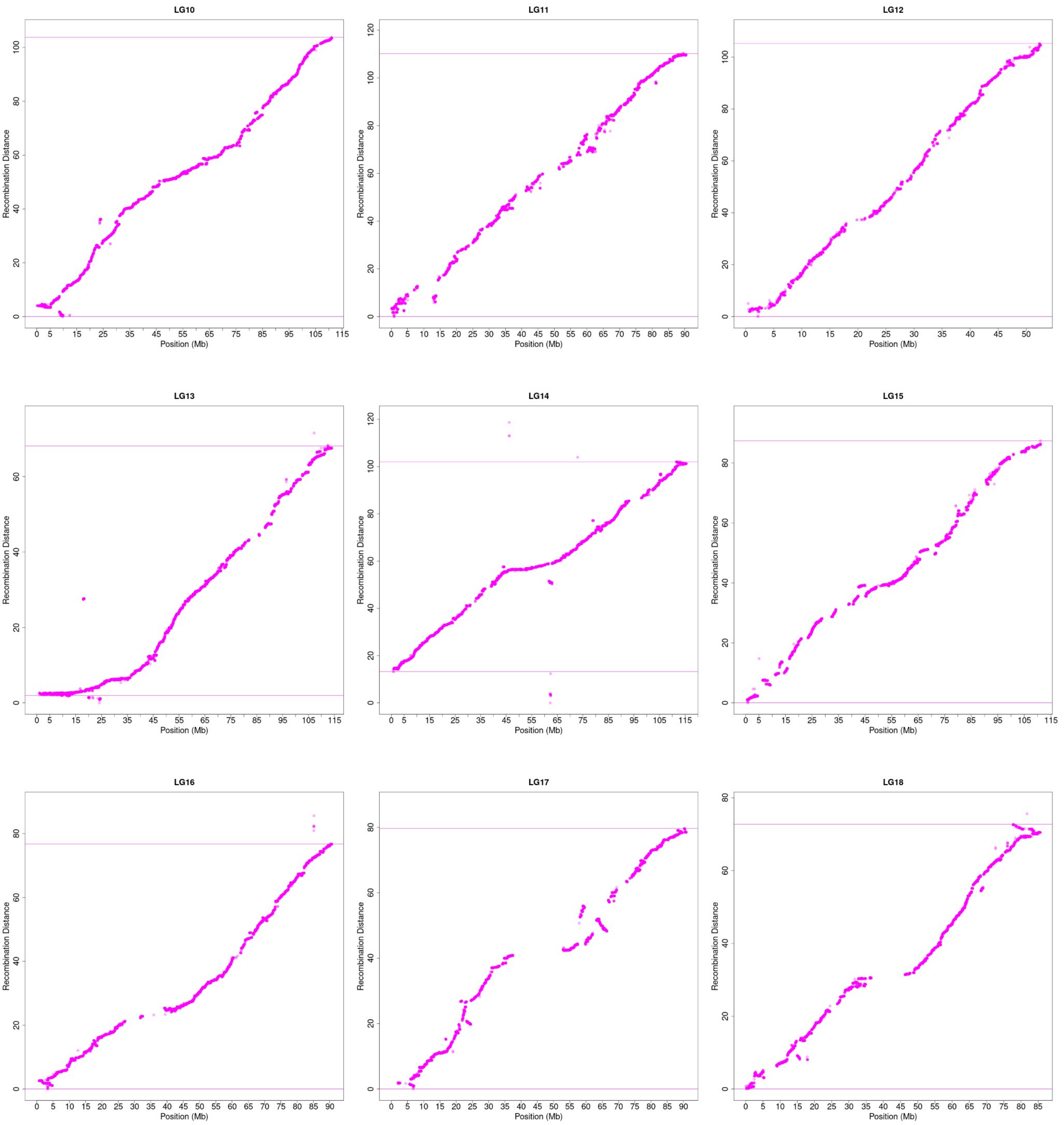


**Figure S5 Continued.**

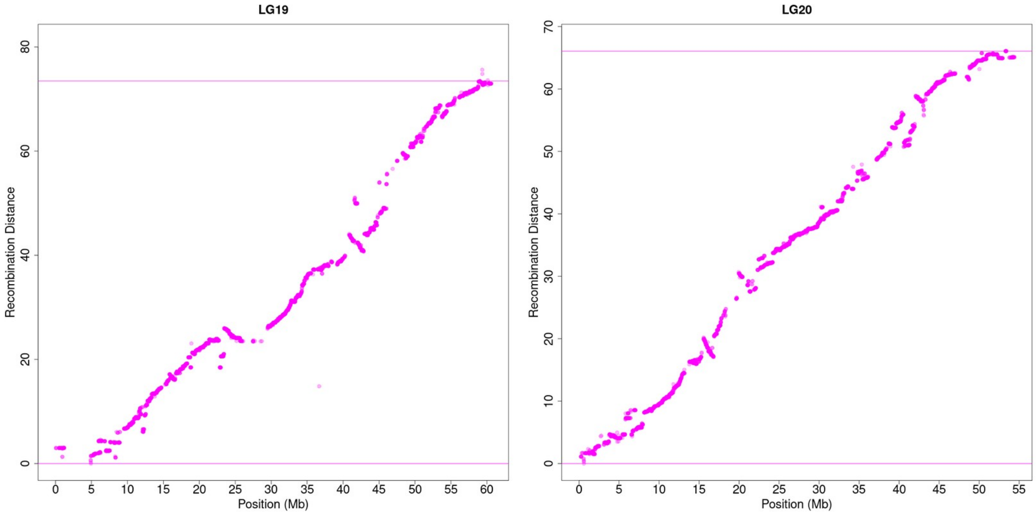


**Figure S5 Continued.**

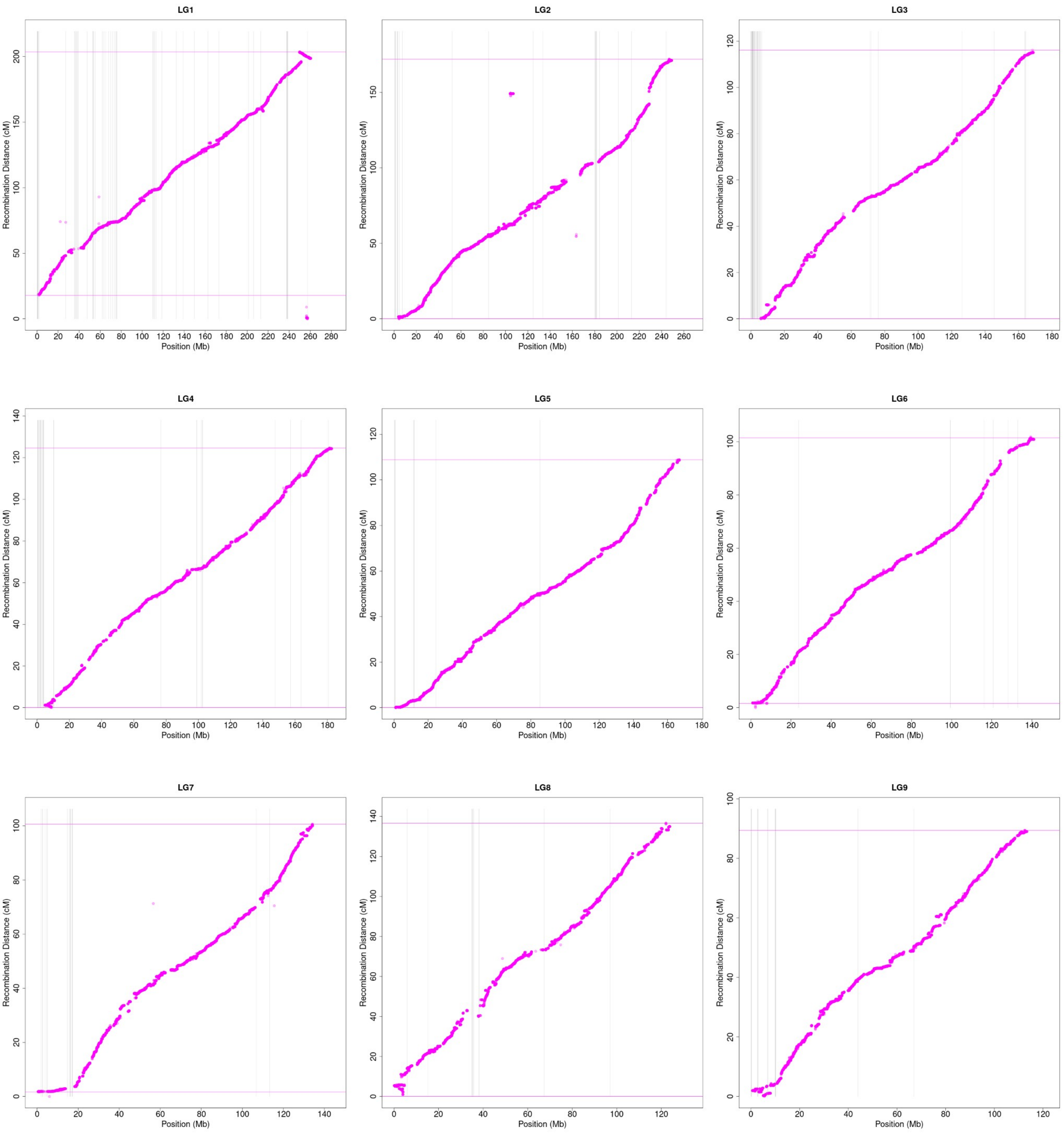


**Figure S6. The order of genetic markers and the distances from a rat genetic map compared to their locations in mRatBN7.2**

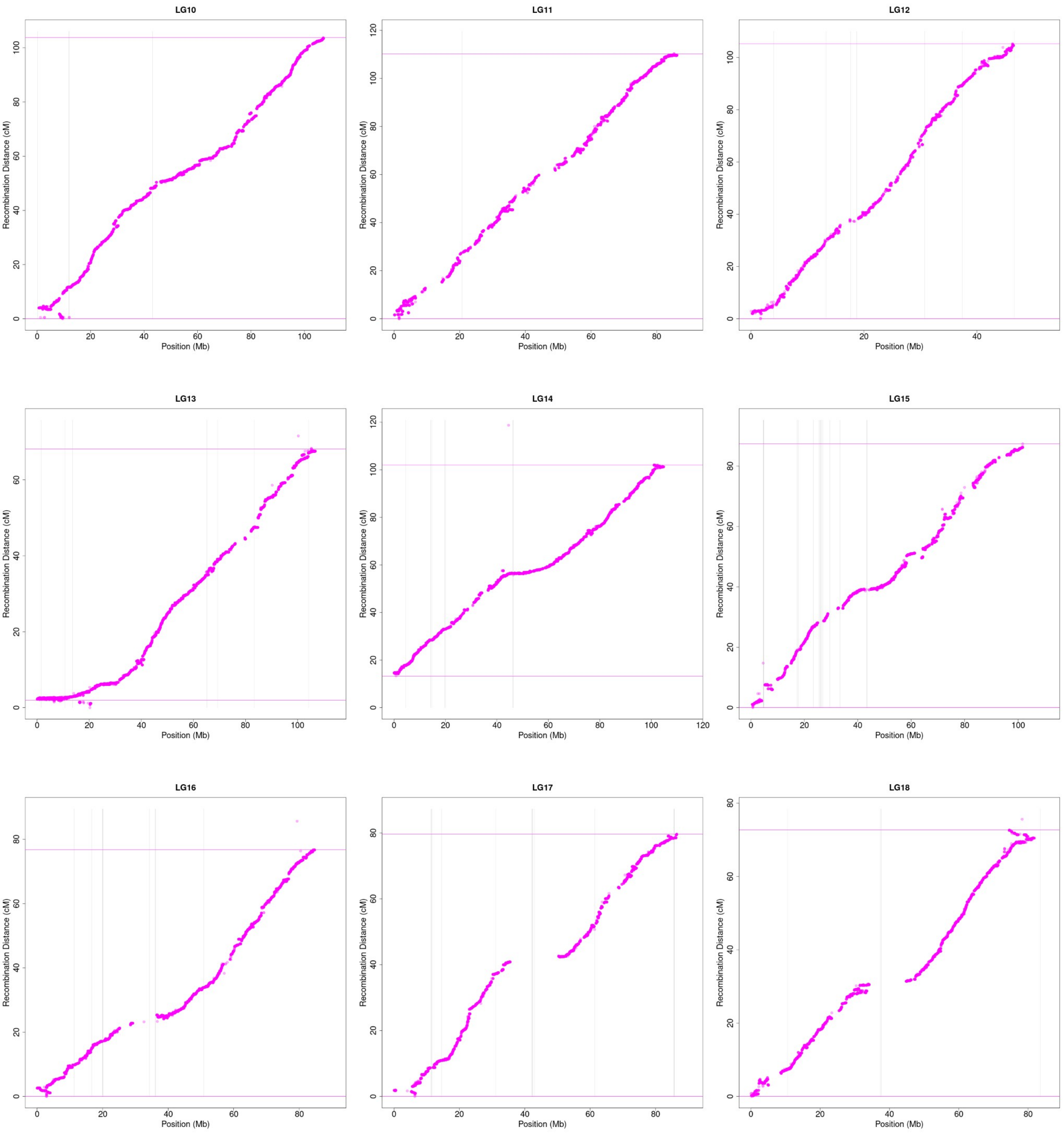


**Figure S6 Continued**

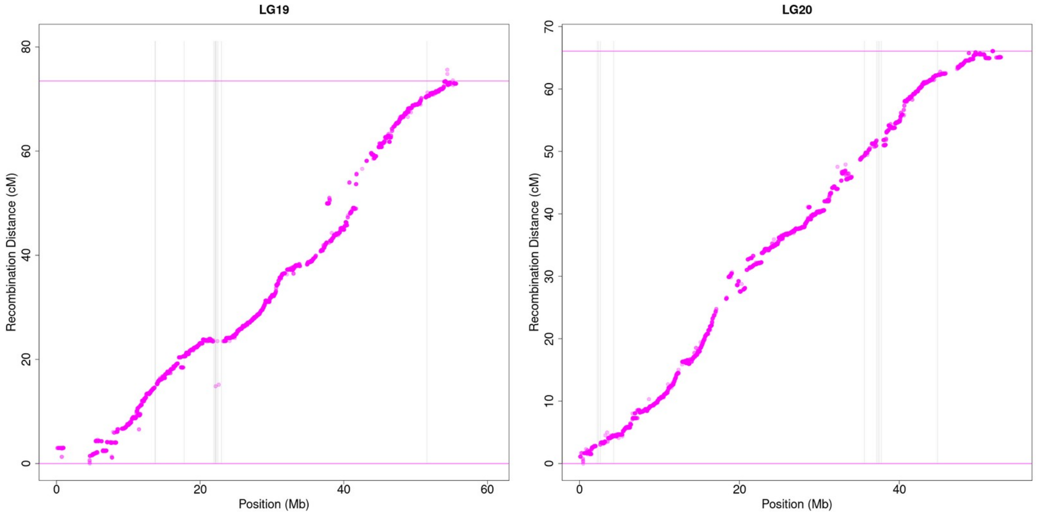


### Figure S6 continued

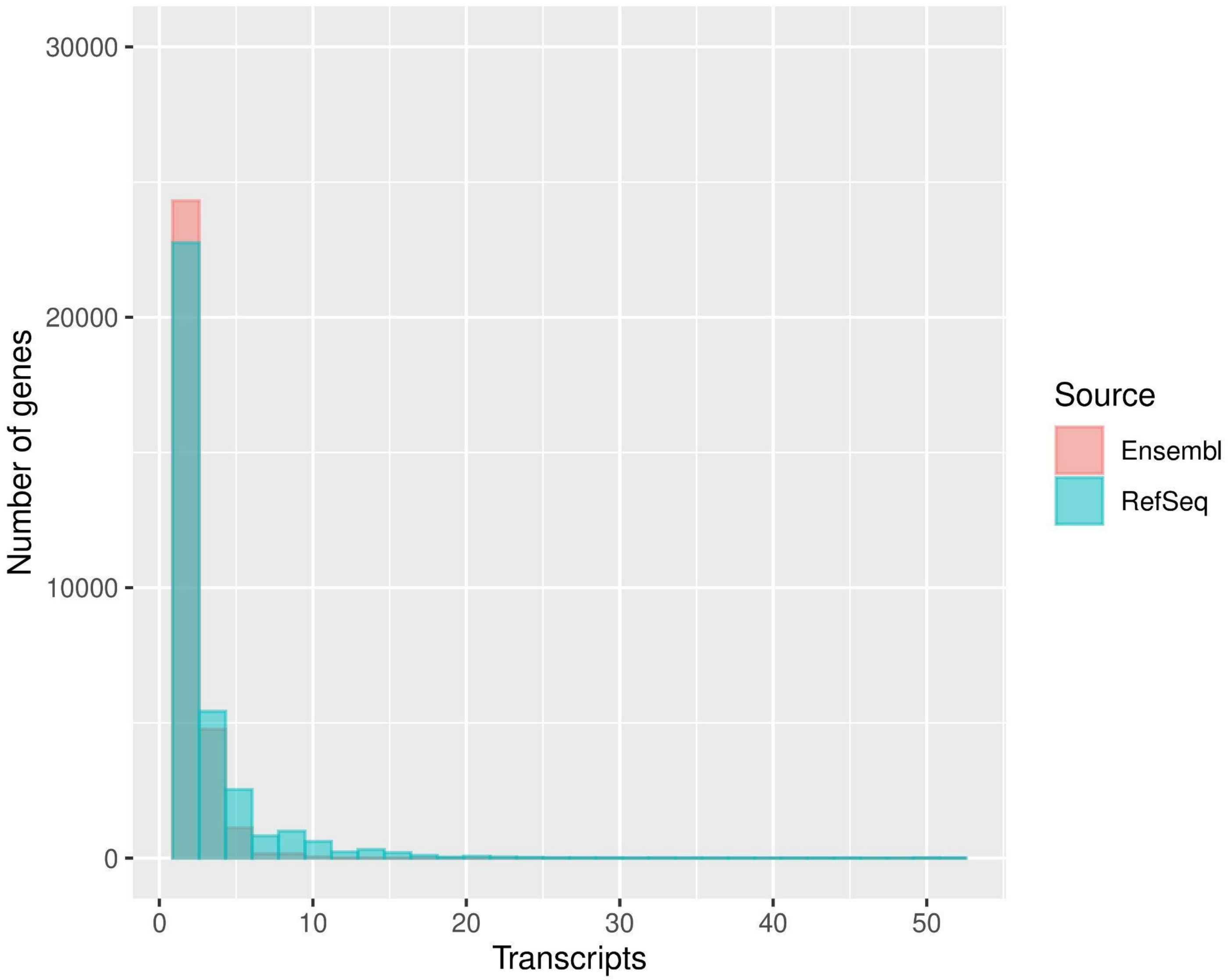


**Figure S7. The number of transcripts annotated by Ensembl and RefSeq.** Refseq annotated a greater number of genes with multiple transcripts than Ensembl. The average number of transcripts per gene was 2.9 for RefSeq and 1.8 for Ensembl.

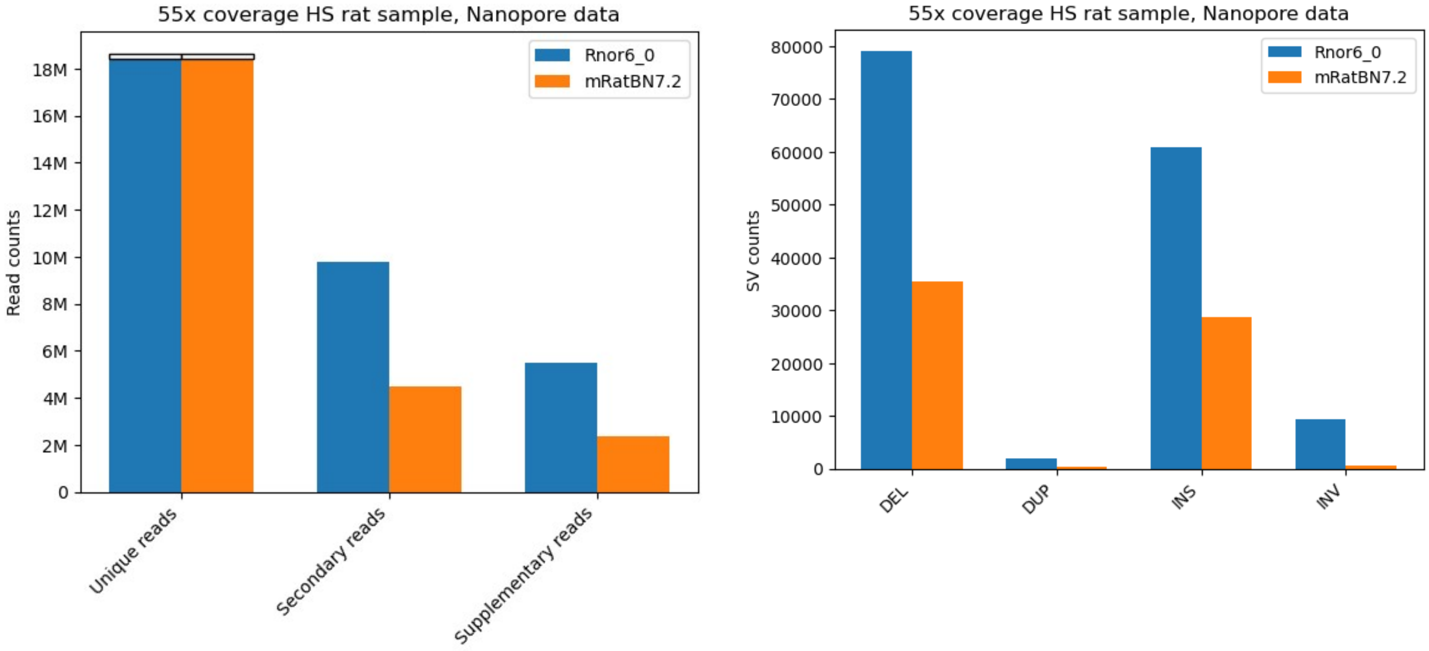


**Figure S8. Mapping results of nanopore data of one HS rat with 55x coverage.**

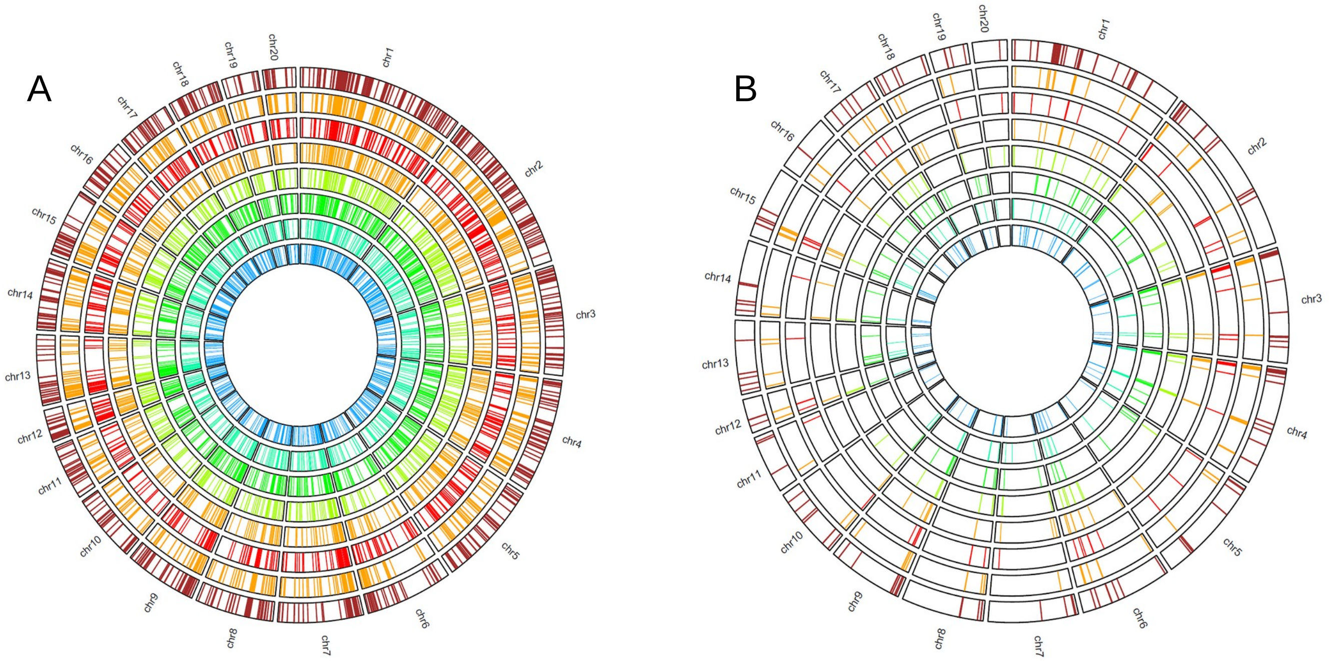


**Figure S9. Locations of large deletions from multiple linked-read samples.** From outer circle to inner circle are the following strains: BN, SHR/Olalpcv, BXH10, BXH8, BXH2, HXB17, HXB2, and HXB21. **A)** Large deletions mapped to Rnor6.0 **B)** Large deletions mapped to mRatBN7.2.

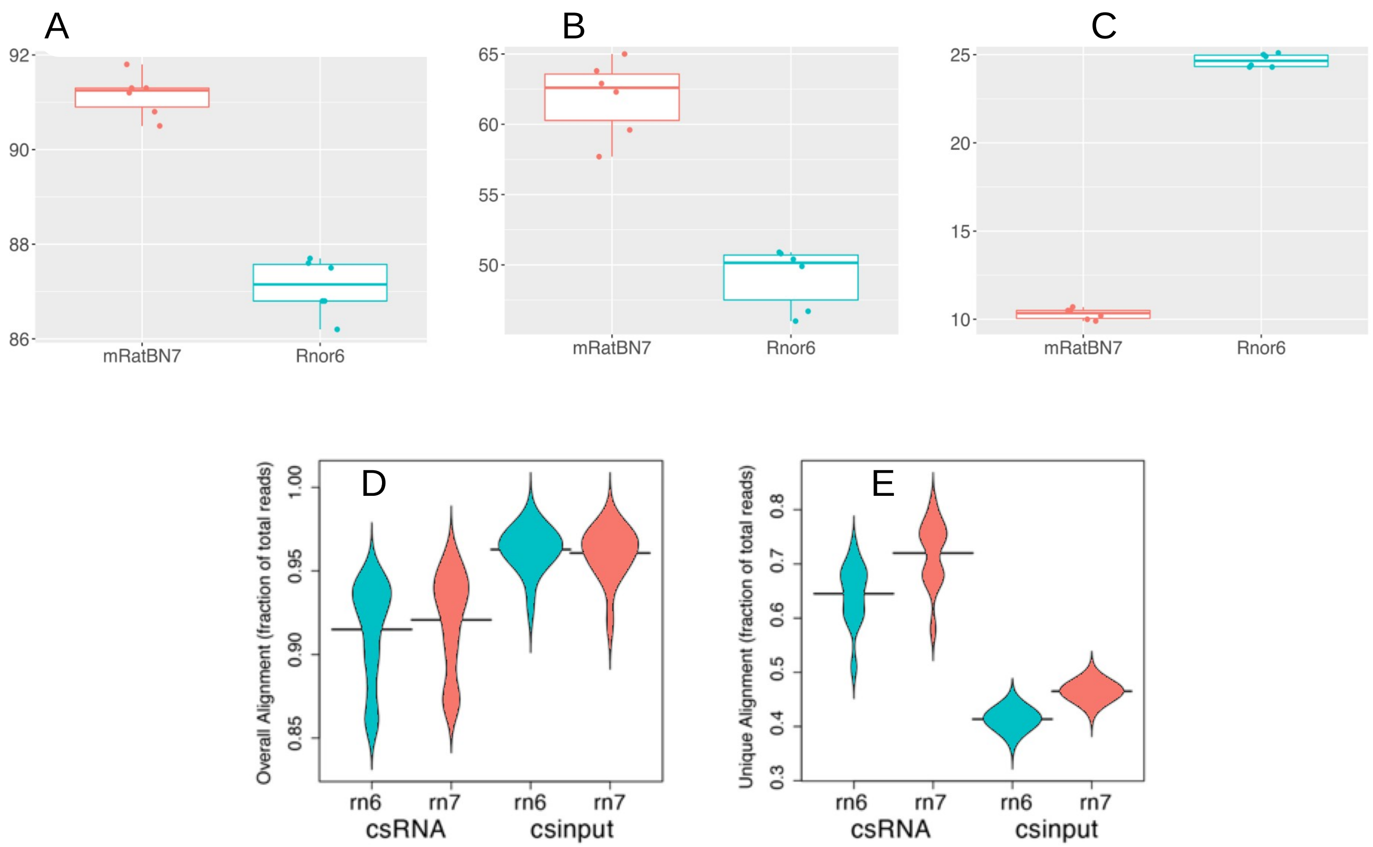


**Figure S10. Mapping metrics of single nuclei RNA-seq and small capped RNA-seq data generated using rat brain tissues.** Percentage of snRNA-seq reads mapped to **A)** the genome, **B)** the transcriptome, and **C)** an intergenic region of the genome for mRatBN7.2 and Rnor_6.0. **D)** Overall mapping rates of csRNA-seq data to two reference genomes. **E)** Unique alignment of csRNA-seq data to two reference genomes.

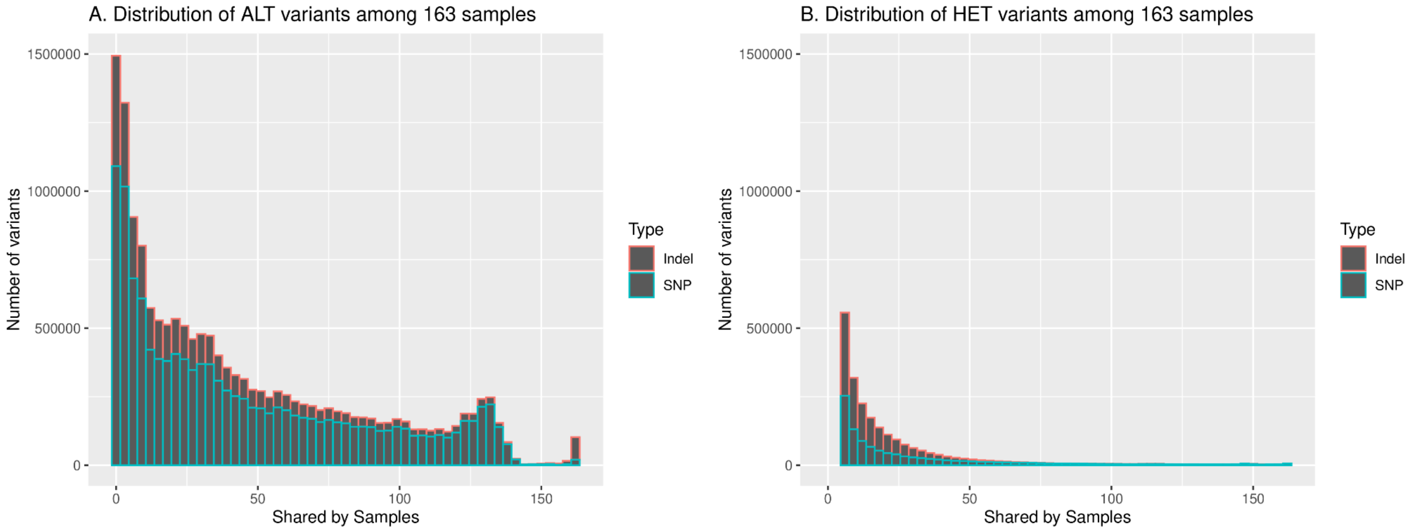


**Figure S11. Distribution of the total number of variants shared by 1 or more samples across all 163 samples** for A) Homozygous SNPs/Indels. The lack of variants shared by approximately 140-155 samples and the distinct peak after 155 samples indicate variants shared by more than 156 samples are likely caused by errors in the reference genome. B) Heterozygous SNPs/Indels. Similar pattern as A can be observed.

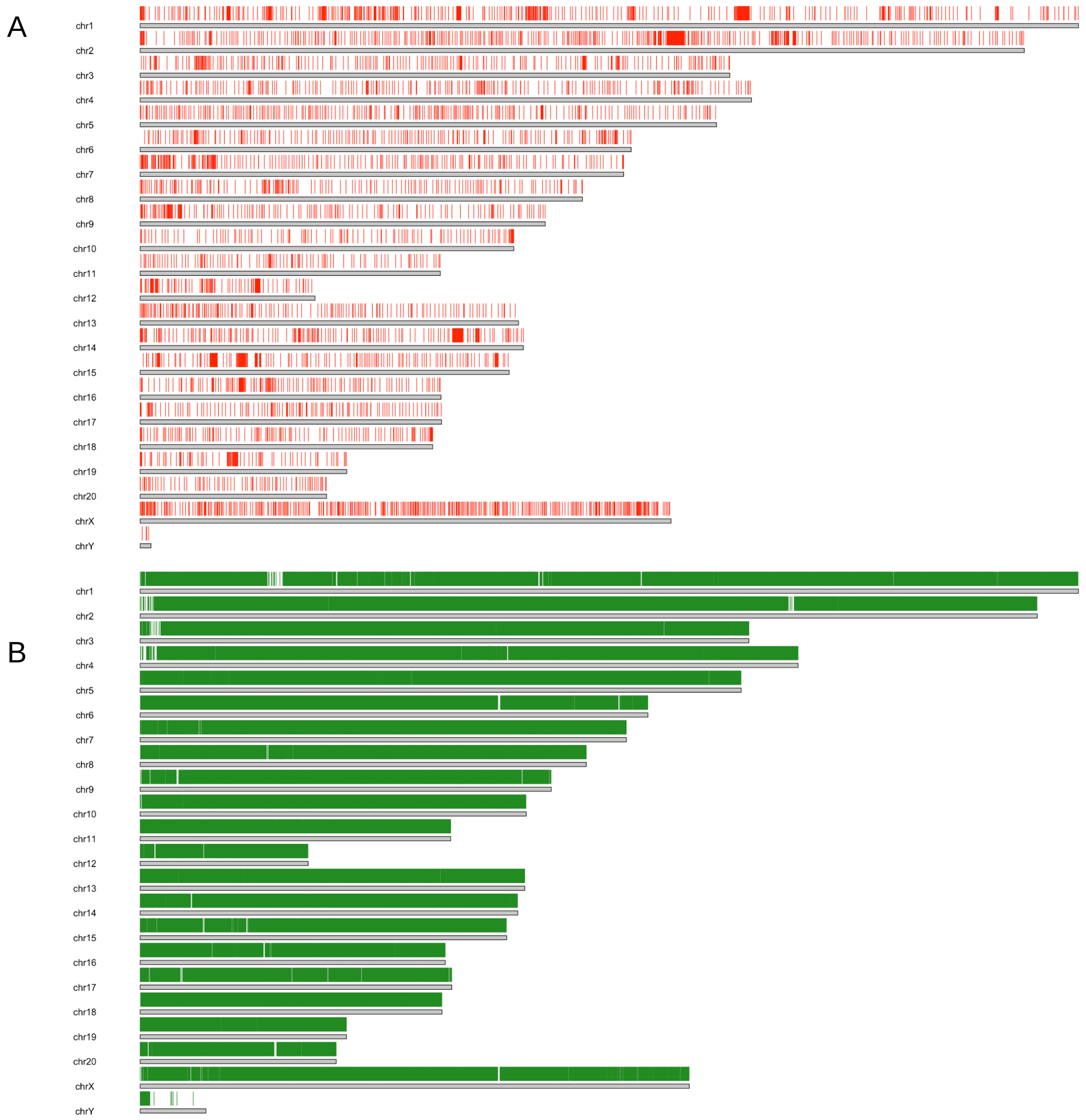


**Figure S12. Simulated liftover analysis: unliftable and lifted site distribution.** An evenly 1000 bp spaced bed file covering Rnor_6.0 is generated and then lifted to mRatBN7.2. Out of the 2,782,023 sites, 92.04% are liftable, 7.96% are not liftable. **A)** The distribution of the unliftable sites are plotted on Rnor_6.0. **B)** The distribution of the lifted sites are plotted on mRatBN7.2. The data are subsampled by 50 for better visualization.

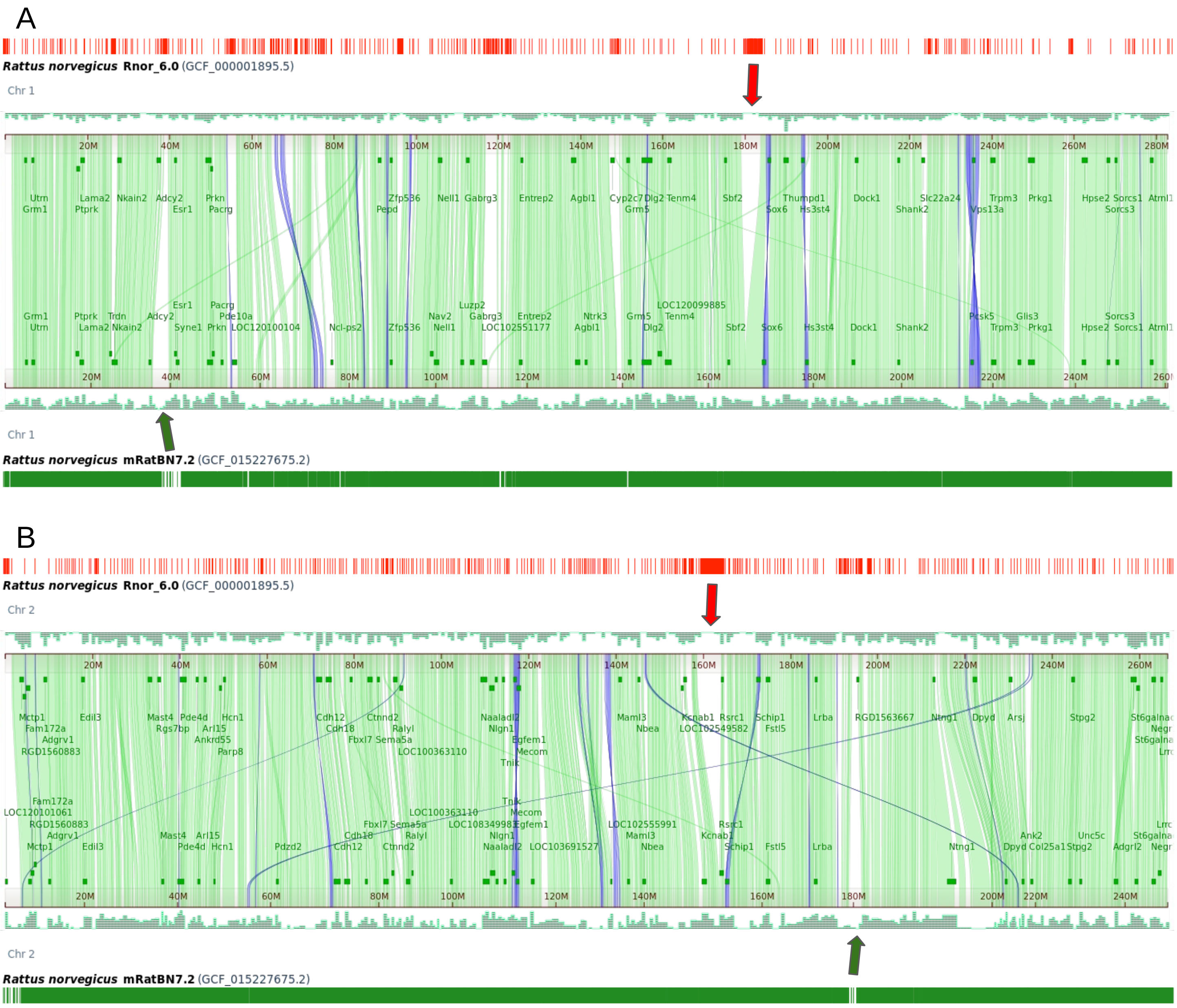


**Figure S13. Simulated liftover analysis and comparative genome view.** Top red track is the unliftable sites distributed on the Rnor_6.0. Bottom green track is the lifted sites distributed on the mRatBN7.2. The middle is the NCBI comparative genome viewer between chr1 of Rnor_6.0 and mRatBN7.2. The red arrow highlights a region that's not liftable in Rnor_6.0. According to the comparative genome view, the region has no corresponding region in mRatBN7.2 (not just chr1). The green arrow highlights a regiono in mRatBN7.2 that has no lifted sites from Rnor_6.0. According to the comparative genome view, this region has matching region in Rnor_6.0. **(A)** Chromosome 1. **(B)** Chromosome 2.

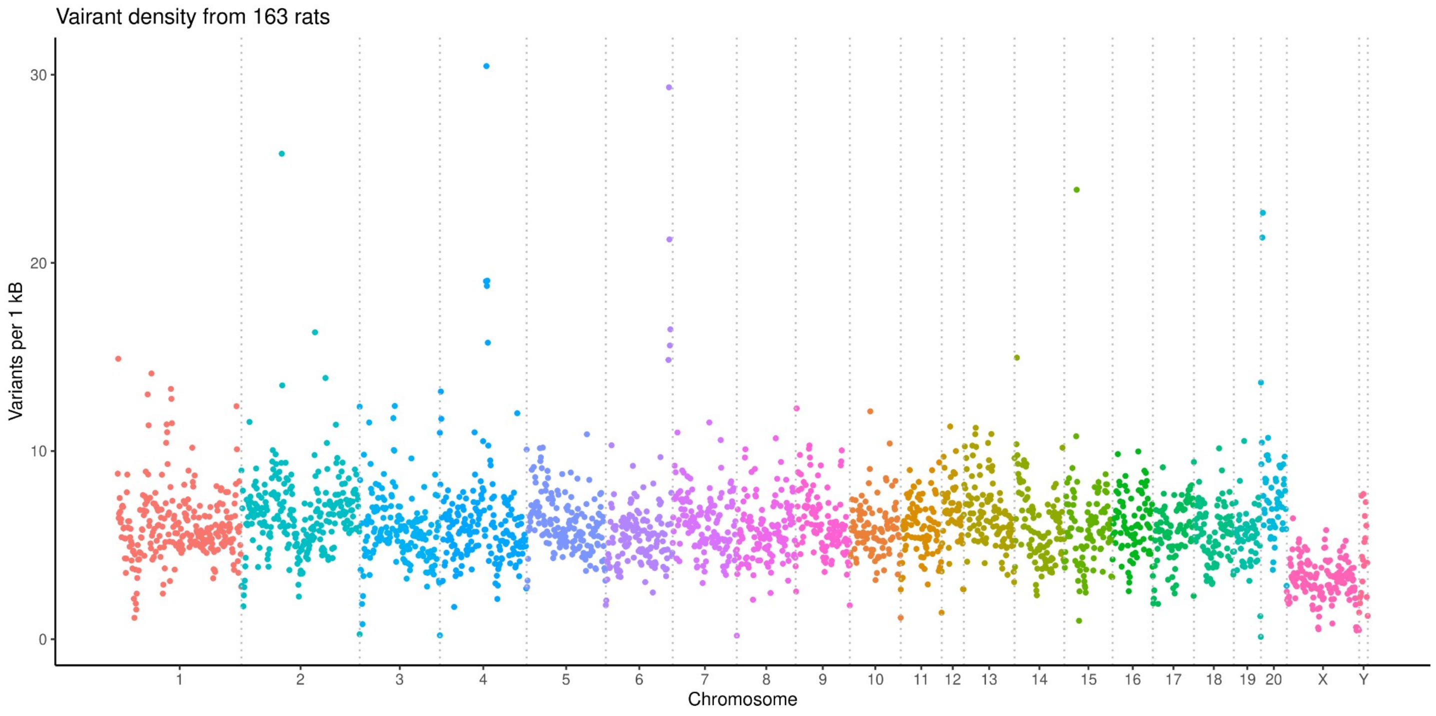


**Figure S14. Density of variants across the genome in a collection of 163 rat WGS samples**

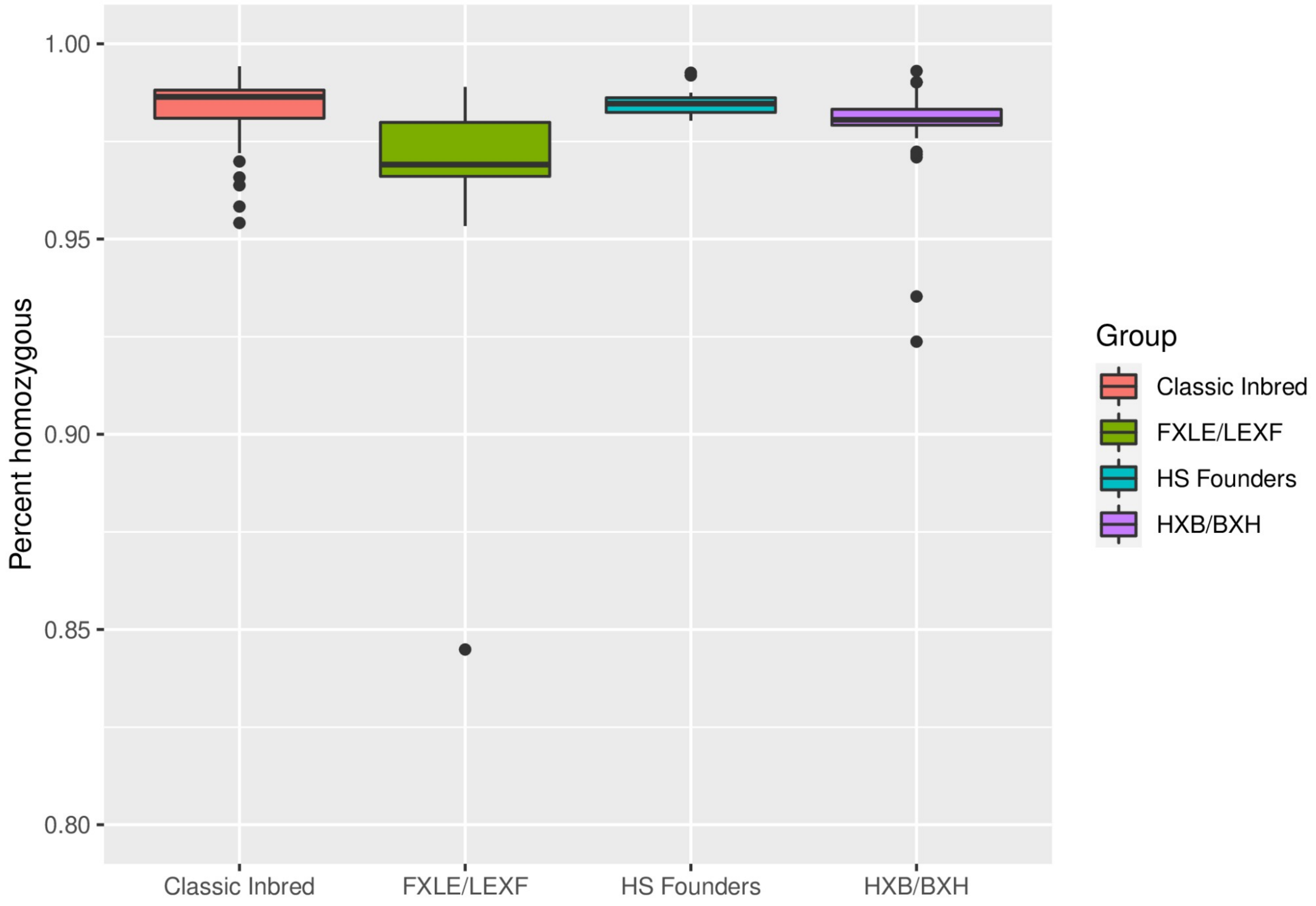


**Figure S15. Homozygosity of inbred strains.** Variants in most sample were homozygous, confirming the inbred nature of most strains. A few exceptions were noted. For example, 15.6% of the variants from FXLE24 were heterozygous. Additionally, BXH2, which we sequenced two samples, has ~7.7% heterozygous variants.

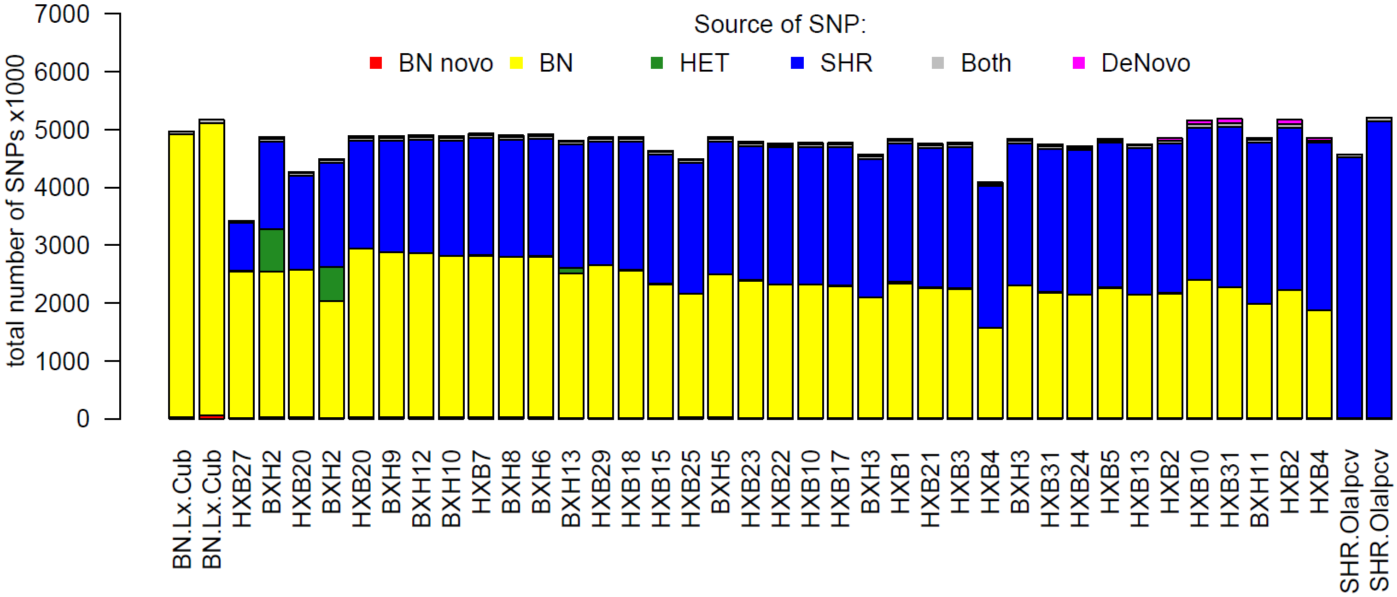


**Figure S16. The majority of the variants in HXB/BXH RI panel are inherited from the parental strains.** De Novo Mutations on BN.Lx (Red) Mutations originating from BN (Yellow), Heterozygous (Green), Mutations originating from SHR (Blue), SNPs found on both BN and SHR (Grey), De Novo SNPs from neither parental strains (Pink).

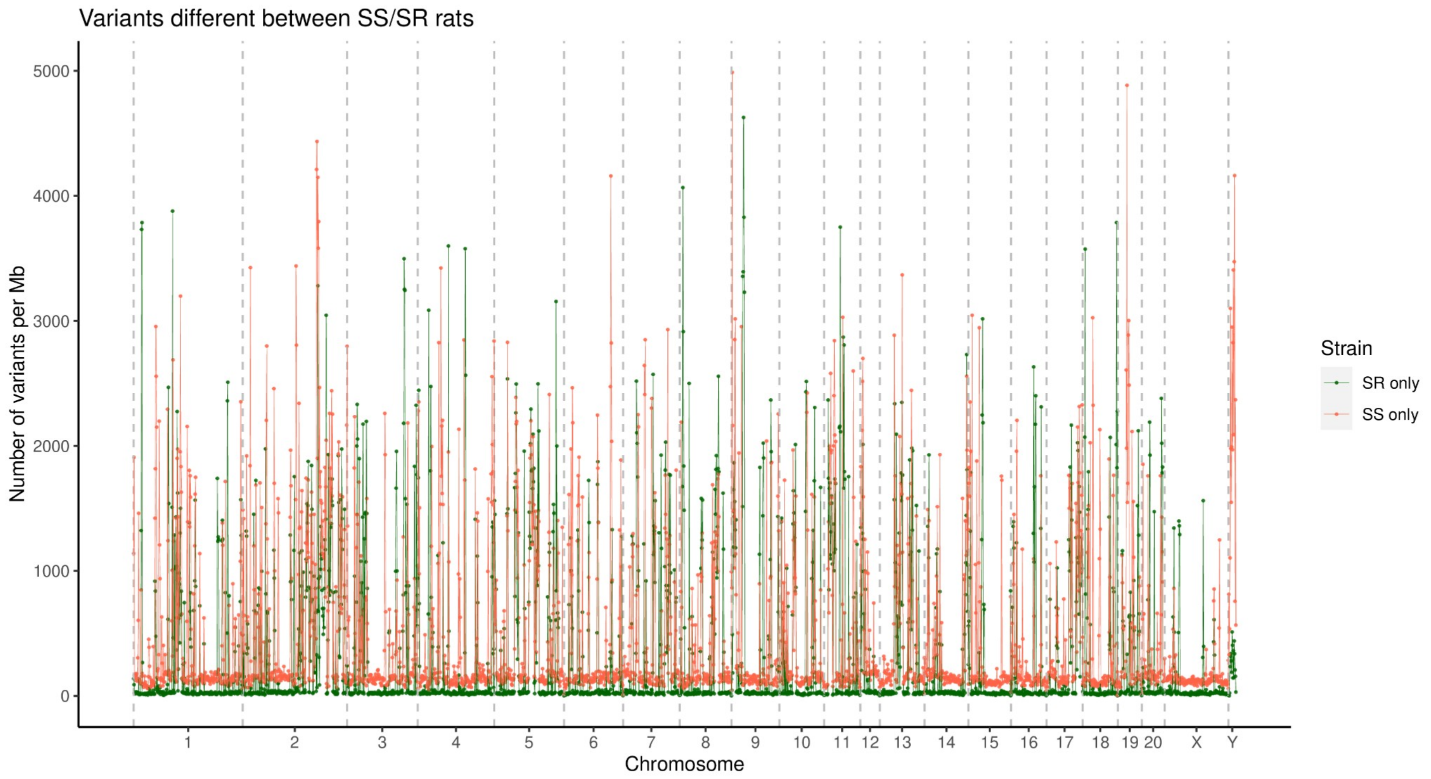


**Figure S17. Distribution of variants different between SS and SR rats.** Total number of variants unique to either SS or SR rats across the genome. The SS strain contains more unique variants than SR.

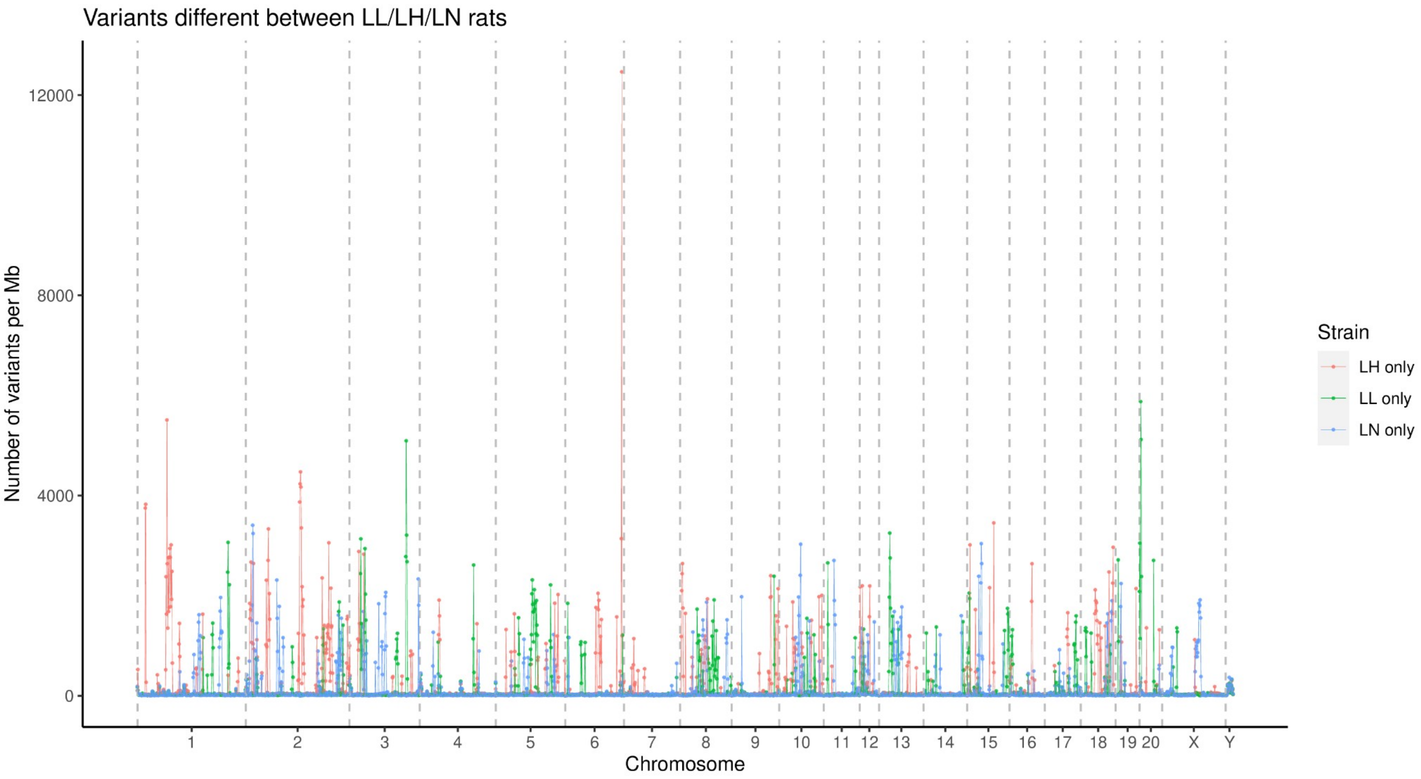


**Figure S18 Distribution of variants unique to LL/LH/LN rats.** Strain-specific variants tend to cluster by strain.

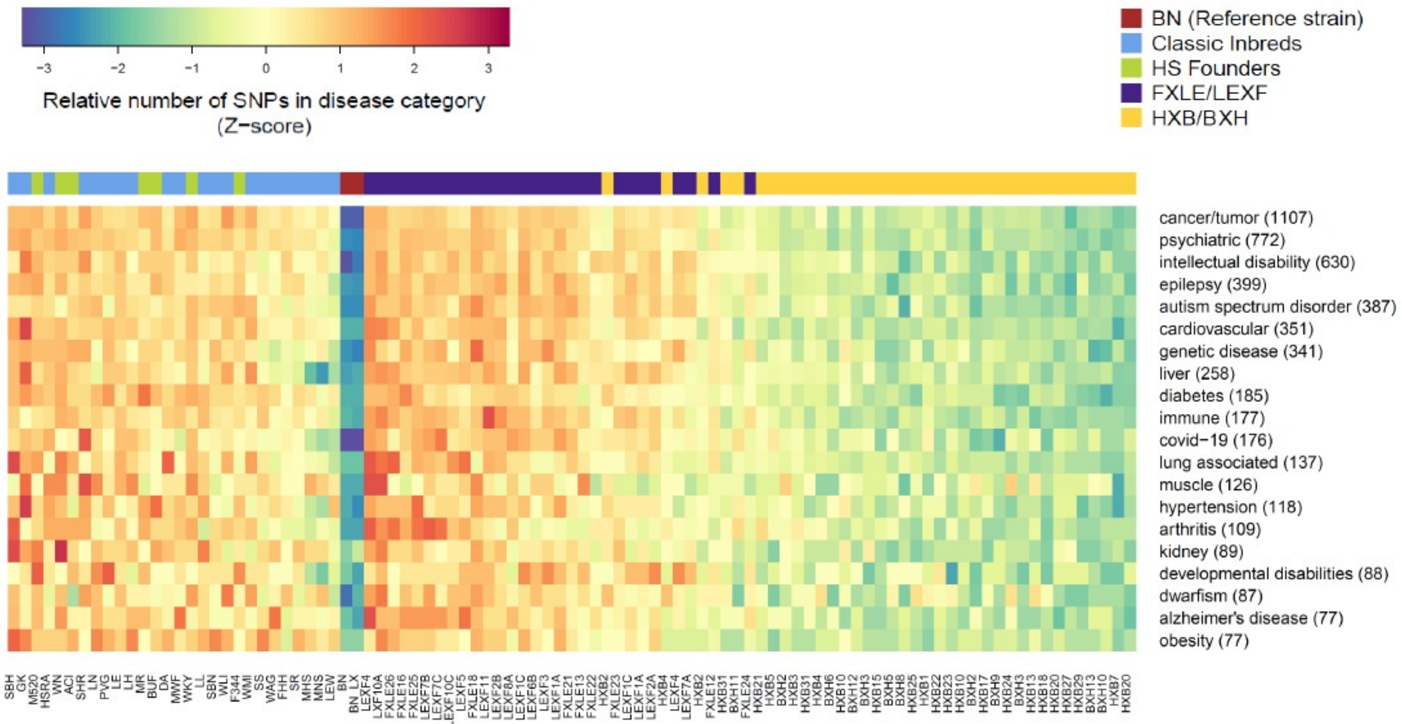


**Figure S20. Overview of disease ontology annotation of variants in RatCollection.** Heat maps shows relative number of SNPs in proximity of genes associated per disease category within the reference, classic inbreds, HS progenitors and recombinant inbred panels.

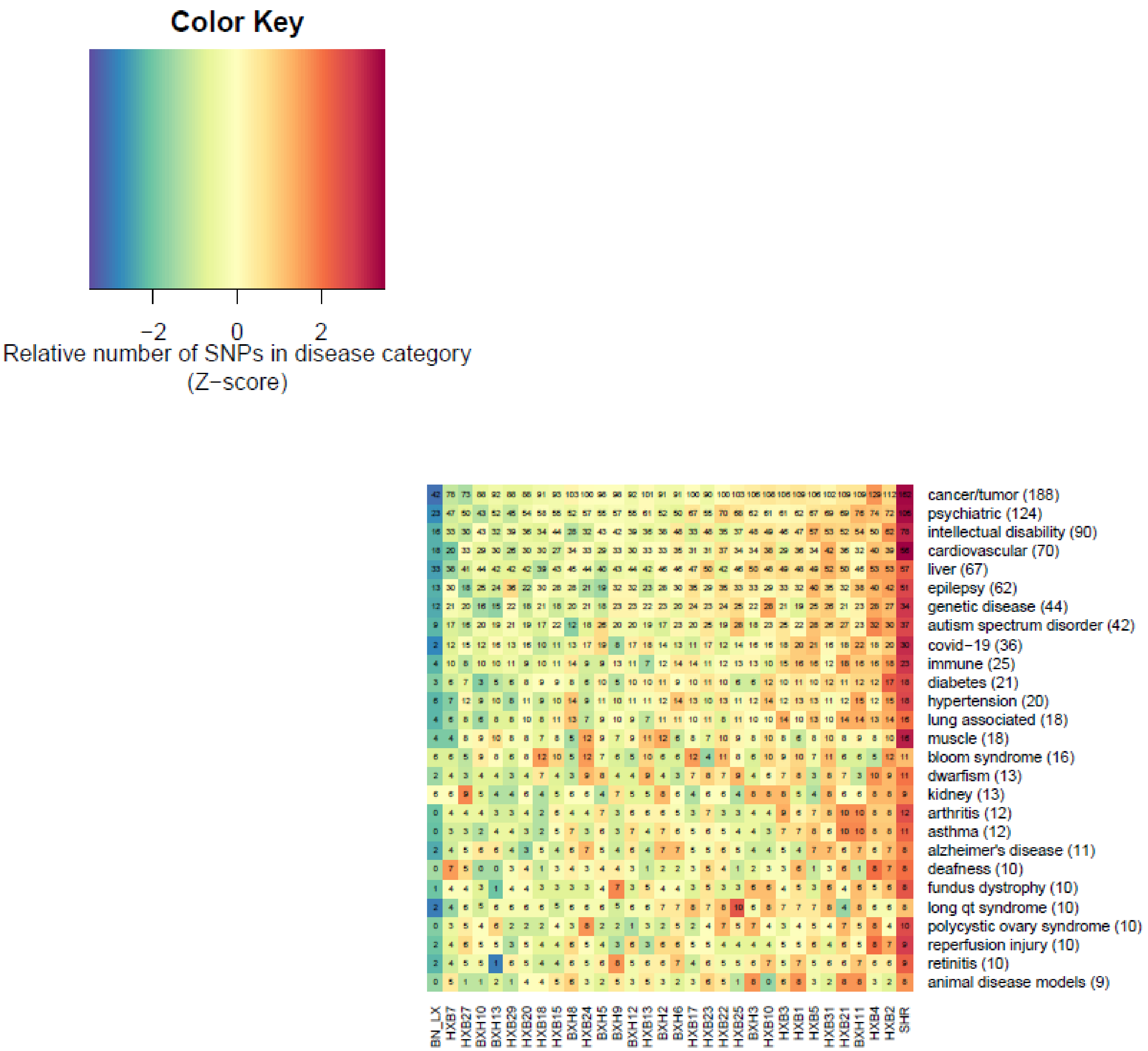


**Figure S21. Disease ontology of genetic variants found in the HXB/BXH RI panel.** Numbers within the grid show the absolute number of SNPs with high impact on genes within the disease annotation. Numbers following the disease name shows the total number of annotated genes with at least 1 high impact variant within the panel.

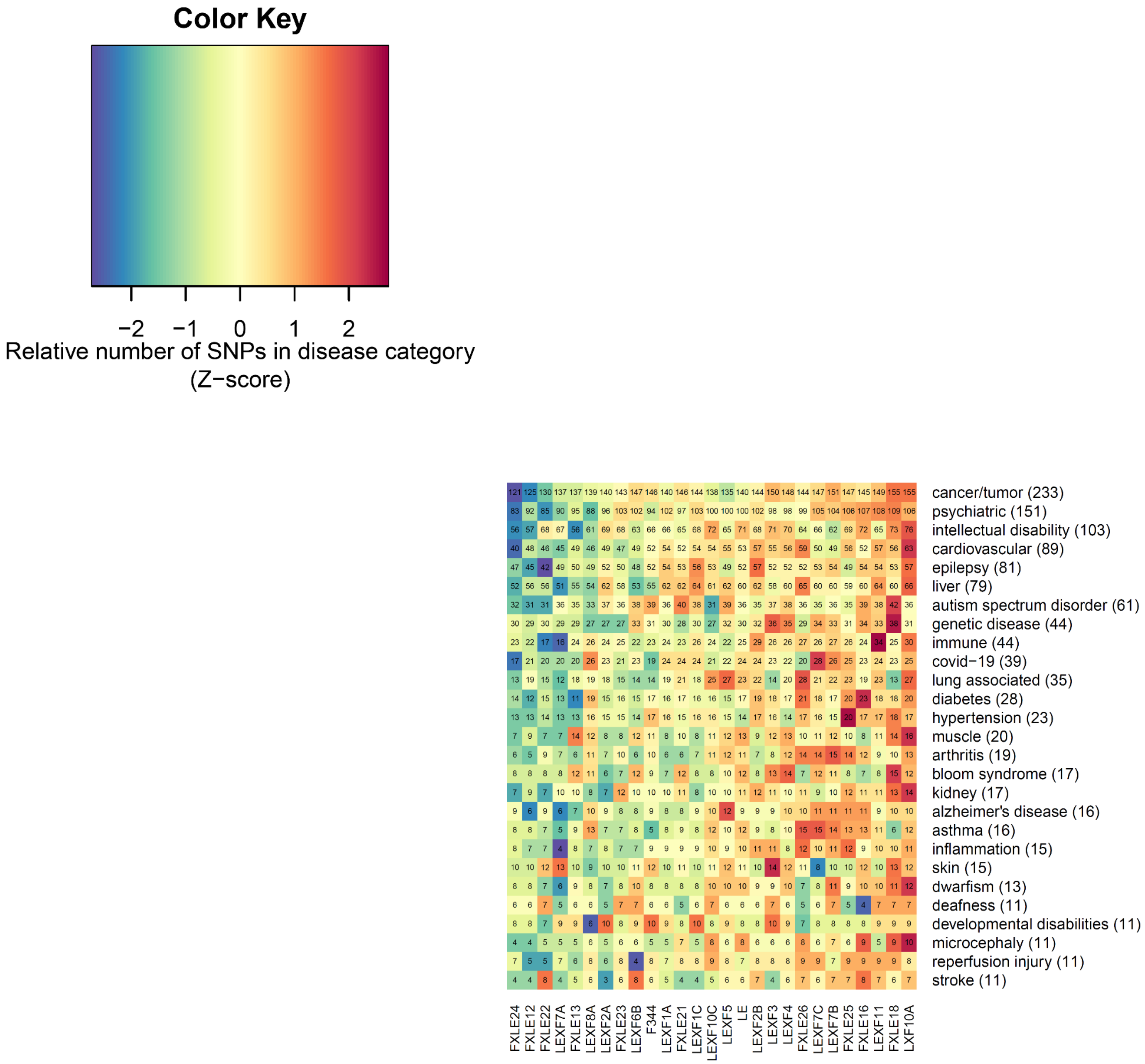


**Figure S22. Disease ontology annotation of variants in the LEXF/FXLE RI panel.** Numbers within the grid show the absolute number of SNPs with high impact on genes within the disease annotation. Numbers following the disease name shows the total number of annotated genes with at least 1 high impact variant within the panel.
